# Comparative Assessment of the Utility of Co-Folding and Docking for Small-Molecule Drug Design

**DOI:** 10.64898/2025.12.09.693161

**Authors:** Ute F. Röhrig, Marine Mathieu-Bugnon, Vincent Zoete

## Abstract

Deep-learning based co-folding methods predict the structures of proteins interacting with metal ions, small molecules, nucleic acids, peptides, and other proteins. One of their main objectives is their application for drug design, predicting the structure of small-molecule ligand/protein complexes. It has been shown that at present these models memorize ligand poses from the training data but do not generalize effectively to novel complexes and lack in the adherence to physical and chemical principles. Here, we use the recently introduced Runs N’ Poses benchmark set of 2,600 protein-ligand systems annotated by their similarity to the training data, to show that the physics-based docking algorithms Attracting Cavities and AutoDock Vina outperform co-folding methods for novel ligands and novel binding pockets. In addition to predicting ligand poses and at variance with co-folding methods, they provide a physical rationale on why a ligand binds (or does not bind) and insight into experimental structural model deficiencies.

## 1 Introduction

The process of developing small molecules as therapeutic agents for unmet medical needs heavily relies on computational methods. At the hit finding, hit-to-lead, and lead optimization stages, computational structure-based drug design aims to predict the three-dimensional structure and interactions between a biological macromolecular target and a potential small-molecule modulator to inform experimental approaches. Predicted interactions are essential for supporting drug design projects, because they allow for rational chemical modifications to improve ligand properties such as affinity or specificity. The core of drug discovery is the generation of novel compounds for novel targets, allowing to secure intellectual property and a competitive advantage.

Deep-learning based co-folding models such as AlphaFold 3,^1^ Umol,^2^ RoseTTAFold All-Atom,^3^ Chai,^4^ Boltz,^5–7^ Protenix,^8^ and NeuralPLexer^9^ show great promise in predicting the three-dimensional structures of biomolecular complexes from amino acid sequences and chemical ligand structures alone. AlphaFold 3 has been advertised as providing “far greater accuracy for protein–ligand interactions compared with stat-of-the-art docking tools.”^1^ However, it has subsequently been shown that at present co-folding models memorize ligand poses from the training data but do not generalize effectively to novel complexes^10^ and lack in the adherence to physical and chemical principles.^11^

Traditional small-molecule docking algorithms on the other hand use three-dimensional target coordinates and (generally) the location of a binding pocket as input, and predict the binding mode of small molecules to this site.^12^ They include a sampling and a scoring step, where the scoring function can be empirical, knowledge-based, force-field based, or a mixture of these categories. The main shortcoming of docking is the static approach to the dynamic ligand-binding process, leading to a reasonable accuracy in predicting binding poses but difficulties in predicting binding affinities and in separating binders from non-binders. However, they have proven their usefulness in numerous drug design projects over the last decades. Since the inception of co-folding methods, these docking algorithms are often called “physics-based”. Independent from the nature of their scoring function, which may be more or less physical, this label refers to the fact that they try to predict binding interactions based on physical principles such as, for example, electrostatic and van der Waals interactions. On the other hand, deep-learning docking algorithms such as DiffDock^13^ also require three-dimensional target coordinates but rely on learned representations for pose generation and scoring.

Co-folding and docking are fundamentally different techniques and therefore difficult to compare. However, as co-folding methods claim to be better at predicting ligand–protein structures, a comparison is warranted. The recently developed Runs N’ Poses (RNP) set provides an optimal starting point for a comparison, as it gives access not only to co-folding results for AlphaFold 3, Chai-1, Protenix, Boltz-1, Boltz-2, and RoseTTAFold All-Atom, but also to wealth of metadata, including similarity measures to both protein binding sites and ligand poses used for model training.^10^ The authors showed that the performance of co-folding methods strongly depends on similarity to the training data and that they generalize poorly to novel complexes.

Here, we first analyze the properties and the quality of the large RNP benchmark set in comparison to the much smaller Astex Diverse^14^ and PDBbind v2016 core (CASF-2016) sets,^15^ which have been widely used for the assessment of drug design tools. Based on the analysis, we define the more balanced and higher quality filtered subset RNP-F (RNP filtered). We then present results for the RNP and RNP-F sets with different physics-based docking algorithms. We deliberately chose to perform re-docking and not cross-docking calculations and to locally dock to a predefined ligand binding site and not blindly on the full protein surface. The first choice was motivated by a better standardization and comparability of docking results, but the provided success rates should be considered an upper bound of what can be expected in a predictive cross-docking setting. The second choice makes sense in a drug design perspective, where a specific ligand binding site is usually targeted and may be predicted beforehand with dedicated pocket prediction algorithms. For docking, we use our algorithm Attracting Cavities (AC)^16–18^ as well as the popular AutoDock Vina (Vina)^19,20^ algorithm. Attracting Cavities with its thorough sampling strategy and universal CHARMM force-field^21,22^ based scoring function with implicit solvation^23^ is freely accessible through the SwissDock webserver^24^ and has been shown to compare favorably to other docking approaches.^17^ We also provide results for the Vina-derived algorithms Smina^25^ with the Vinardo scoring function,^26^ a simplified version of the Vina scoring function, as well as GNINA,^27^ which uses a convolutional neural network scoring function. Finally, we compare the docking results to the AlphaFold 3 binding pose predictions provided with the RNP set.^10^ We limit comparison to AlphaFold 3 as all co-folding methods were shown to behave similarly and in order to keep as many cases as possible in the common subset solved with all methods. We provide all docking results including input files and annotations at https://doi.org/10.5281/zenodo.17865538.

## 2 Methods

### 2.1 Data Sets

The Runs N’ Poses set (github.com/plinder-org/runs-n-poses)^10^ is composed of 2,600 protein-ligand systems with 3,047 “proper ligands” from the protein data bank (PDB)^28,29^ released between the training cutoff date for the co-folding methods (October 1, 2021) and before January 9, 2025. Proper ligands were defined by PLINDER^30^ as ligands not labeled as ions or artifacts. Binding site similarity was quantified by the “pocket coverage”, i.e. the percentage of residues within 6 Å of the ligand in the query system which align with residues within 6 Å of the ligand in the target system (regardless of amino acid identity). Ligand pose similarity was quantified by the SuCOS score (github.com/susanhleung/SuCOS),^31^ which measures the overlap of ligand poses in volume and placement of chemical features. Overall similarity was defined as the product of these two measures, divided by 100 to obtain a parameter in the interval 0–100. For 2,812 proper ligand-protein systems, the ligand and pocket similarity scores were provided. This set was divided into 921 clusters based on graph community clustering at ligand/pocket similarity > 50.^10^

For comparison with well-established docking benchmark sets we used the Astex Diverse set^14^ of 85 complexes and the PDBbind core set v2016,^15^ consisting of 57 target clusters with 5 complexes each for a total of 285 ligand–protein complexes.

### 2.2 Structure Preparation for Docking

We downloaded all mmCIF files corresponding to the set of 2,812 proper and annonated ligand-protein systems from the PDB^28^ (2,488 different PDB IDs). For each system, we selected the ligand of interest as defined in the RNP set by its _atom_site.label_asym_id. We used ChimeraX^32,33^ to delete the solvent, create all copies of the system which make crystal contacts at a distance 4.5 Å or less, keep only chains at a distance below 8 Å to the ligand, delete cofactors at a distance above 8 Å to the ligand, delete crystallization agents, determine the center of mass of the ligand, add hydrogens to the ligand, save the ligand in mol2 format, remove uncoordinated ions, remove atoms with alternate locations, complete missing sidechains, and delete the ligand of interest before saving the target structure in mmCIF format. This process was successful for 2,806 target and ligand structures out of 2,812.

For 2,611 of these ligand-protein systems, we could generate the necessary structure, topology, and parameter files for CHARMM/Attracting Cavities using the latest versions of SwissParam for all ligands^24^ and CHARMMER, our in-house scripts to prepare CHARMM^21^ input files, which uses the mmCIF parser from PDBeCIF.^34^ The 195 failures (7%) were mainly due to missing ligand or cofactor parameters, for example for boron-containing compounds or polymeric compounds, and to missing co-factor atoms in the PDB model. For each system, we generated randomized ligand conformations with RDKit^35^ and native ligand conformations in sdf and pdb format with Open Babel.^36^

For the same 2,611 ligand-protein systems, we generated target and ligand files for use with AutoDock Vina and its derivatives. We used ChimeraX^32,33^ to convert the initial target files from mmCIF to pdb format, which was then used as input for the prepare_receptor4.py script from MGL tools.^37^ Randomized ligand structures were converted from mol2 to pdbqt format with the prepare_ligand4.py script from MGL tools.

### 2.3 Docking with AC

We used the latest version of AC with improved and accelerated sampling capabilities, which will be described in a forthcoming publication. The scoring function is identical to the one in the latest published version^17^ and the one used on the SwissDock webserver,^38^ so that the conclusions are independent of the employed AC version. All dockings were carried out with the CHARMM36 all-atom additive force field^22,39^ and the CHARMM molecular simulation program,^21^ version 48b1, using the standard AC scoring function (total force-field energy with FACTS^23^ implicit solvation), an initial ligand rotation of 90°, 4 random initial conditions, a rigid protein, and a cubic search box with an edge length of 25 Å centered on the center of mass of the ligand in the corresponding PDB model. The docking calculations provided results for 2,597 out of 2,611 cases (99.5%), belonging to 878 different clusters, the 14 failures mainly being due to issues in the treatment of protein/nucleic acid complexes. We used two different sets of sampling parameters, denoted in the following as AC_norm_ and AC_long_, resulting in maximally 3,072/12,288 initial ligand poses and average CPU times of 45/165 min on a single Intel Core i9 4.7 GHz CPU, respectively. Additionally, we relaxed the PDB pose of each ligand and calculated its score to determine scoring failures (see Sec. 2.5).

### 2.4 Docking with AutoDock Vina and Derivatives

For all dockings, the same search space definition as for AC was used, namely a cubic search box with an edge length of 25 Å centered on the center of mass of the ligand pose in the PDB model. We used AutoDock Vina, version 1.2.3,^19,20^ its Smina fork, version Oct 15, 2019,^25^ with the Vinardo scoring function,^26^ and GNINA, version 1.3.2.^27^ We tried different exhaustivity values (1/8/100) for Vina and the default value of 8 for the other algorithms. Dockings took on average 1 minute with an exhaustivity value of 8 and 15 minutes with an exhaustivity value of 100. Vina and Smina calculations were performed on a single Intel Core i9 4.7 GHz CPU, while GNINA calculations were run on a Nvidia A100 GPU. With each scoring function, we also relaxed the pose of each ligand from its PDB model and calculated its score to determine scoring failures (see Sec. 2.5).

### 2.5 Analysis

In addition to the metadata provided by the authors,^10^ we extracted the X-ray resolution, R-free, and R-work values as well as the ligand B-factors from the PDB^28^ entries of the ligand–protein complexes. Based on the ligand structures at neutral pH generated with the SwissParam approach,^24^ we calculated net ligand charges and recalculated the number of ligand rotatable dihedral angles with RDKit.^35^ For ligand burial we tested both the solvent accessible surface algorithms implemented in ChimeraX^32,33^ and in CHARMM^21^ and found the latter to yield more meaningful results when the two algorithms diverged (see Fig. S1), although its absolute values are higher than intuitively expected. We calculated the electron density score EDIA (electron density support for individual atoms) in its combination for multiple atoms (EDIA_m_)^40^ with the density-fitness tool^41^ from PDB-REDO^42^ for all complexes. We extracted the EDIA_m_ values for each ligand, all residues within 5 Å of the ligand, and all residues within 7.5 Å of the ligand. We also assessed the “molecular beauty” or relevance for drug discovery of all ligands by calculating their OiiSTER map values.^43^

For analysis of docking results, root mean square deviation (RMSD) values to the ligand pose in the PDB model were calculated with spyrmsd,^44^ taking molecular symmetry into account. Local difference distance test of protein-ligand interaction (LDDT-PLI) values^45^ were calculated with OpenStructure.^46^ For AlphaFold 3, the values provided by the authors of the RNP set were used.^10^

Similarity to the training set was defined as in RNP,^10^ using the product of binding pocket coverage^30^ for binding site similarity and the Combined Overlap Score (SuCOS)^31^ for ligand similarity. For comparing co-folding and docking results, success was defined as a combination of a ligand pose RMSD ≤ 2 Å and a LDDT-PLI > 0.8. For analyzing docking results alone, we used different ligand RMSD cutoff values (1.0, 1.5, 2.0 Å) as in our earlier works.^17^ As the protein structure is rigid, the LDDT-PLI is assumed to be correct at low RMSD values. The validity of this assumption is demonstrated in Fig. S2, showing that almost all docking poses with a RMSD below 2 Å have a LDDT-PLI above 0.65.

Scoring failures occur when a scoring function does not attribute the best score to the ligand binding pose defined in the PDB model. They are more likely to be detected with extensive sampling and can be divided into three classes according to their underlying cause: an issue in (1) the scoring function, (2) the structure preparation, or (3) the structural model in the PDB.^17^ To determine scoring failures, we relaxed the PDB binding modes and calculated their score with each scoring function. If we obtained any docking pose with a lower score and a RMSD > 2 Å with respect to the ligand binding mode in the PDB model, a scoring failure was defined.

Logistic regression models for success rates were created with the python modules scikit-learn^47^ and statsmodels,^48^ using the following 5 parameters: ligand pose similarity (ligand SuCOS), binding pocket similarity (pocket coverage), ligand burial, ligand flexibility (number of rotatable dihedrals), and ligand density fit (EDIA_m_). All variables were min-max scaled to the range [0, 1] before regression, removing a single outlier with more than 25 rotatable dihedrals.

## 3 Results and discussion

### 3.1 Analysis and Comparison of Benchmark Sets

We first assessed and compared the properties and the quality of the RNP benchmark set to the Astex Diverse^14^ and the PDBbind sets.^15^ The Astex Diverse set consists of 85 manually selected, pharmaceutically or agrochemically interesting and diverse complexes with high-quality electron density support of ligand poses. The set was developed in 2006 and has been used extensively for the benchmarking of different docking approaches.^49–54^ The PDBbind core set v2016^15^ consists of 57 target clusters with 5 complexes each for a total of 285 ligand–protein complexes. The set was designed for validating docking and scoring methods and is provided with curated starting structures including hydrogen atoms for proteins and ligands on the PDBbind resource (www.pdbbind.org.cd).^55^ In the past, we found a number of errors and problems in this set and provided our own manually curated structures on Zenodo.^56^

Our analysis shows that RNP contains complexes with a broader range of X-ray resolutions and R-free values than both Astex and PDBbind sets, and that ligands are larger and more flexible (Fig. 1). Ligands are on average more solvent exposed than in the Astex set but less than in the PDBbind set, and there are more ligands with a large negative charge. These observations are unproblematic but likely to lead to lower success rates for physics-based docking tools on the RNP set than on the Astex and PDBbind sets.

**Figure 1.**
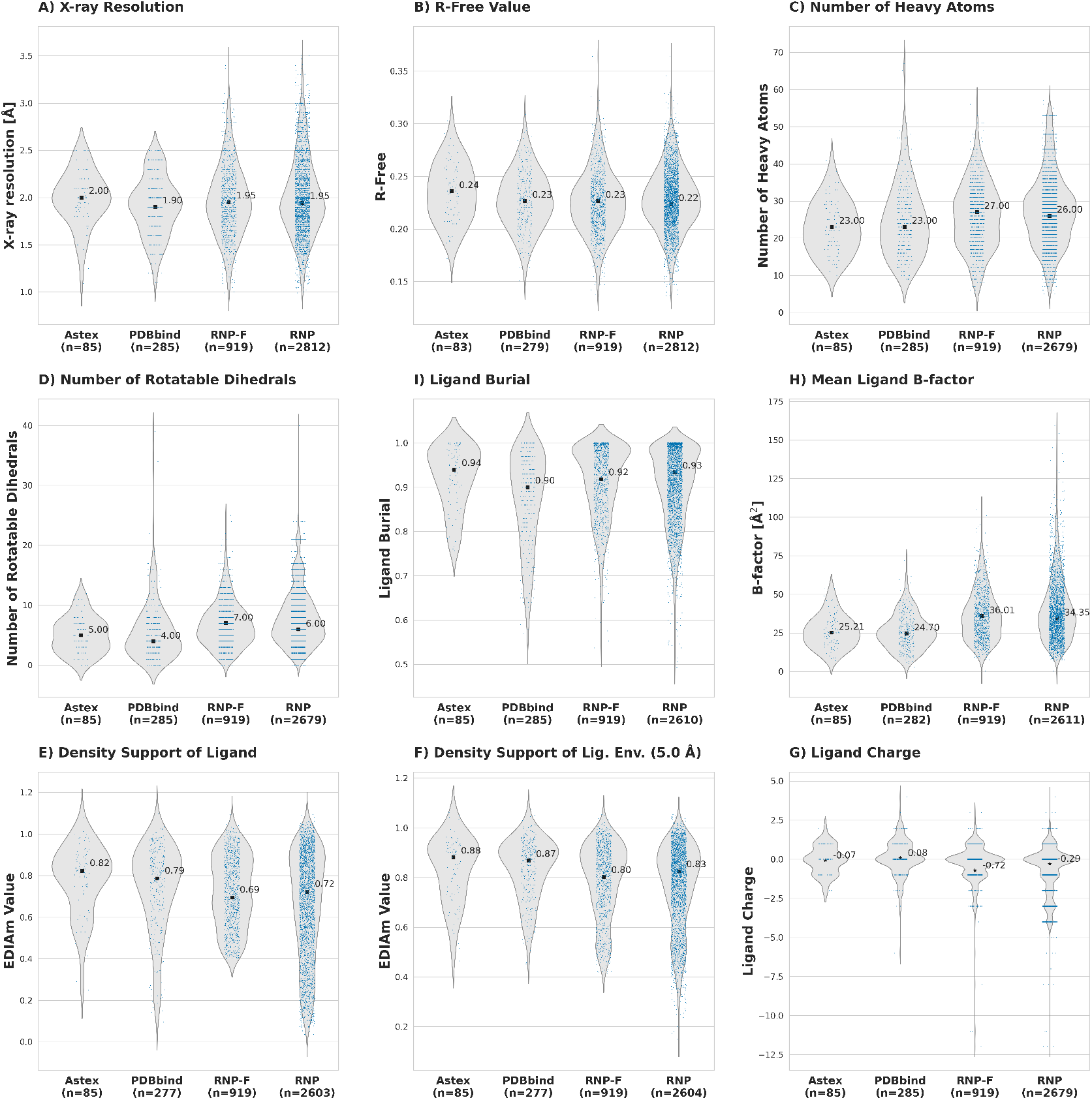
Characteristics of Benchmark Sets. Median/mean values are indicated by black squares/stars, respectively, and annotated. Data shown for Astex, PDBbind, RNP, and RNP-F sets.

However, the RNP also contains a large proportion of complexes, where the ligand and the residues in its immediate environment (within 5 Å) are ill-defined by the electron density, as calculated by their EDIA_m_ scores.^40^ An EDIA_m_ score above 0.8 implies that the molecule is well covered with electron density, a value between 0.4 and 0.8 implies minor inconsistencies with the electron density fit, and a value below 0.4 implies substantial inconsistencies.^40^ 19% of cases in the RNP have a ligand with an EDIA_m_ value below 0.4, compared to 2% in the Astex set and 8% in the PDBbind set. Some examples are shown in Fig. 2B and C, where ligand poses are only partially defined by the electron density. This is problematic, as a poor ligand density fit does not allow to define docking successes and failures, as shown for example in Fig. 2B. Contrarily, in a ligand with a high EDIA_m_ value (Fig. 2A) each atomic position is well defined and can be used as “ground truth”.

**Figure 2.**
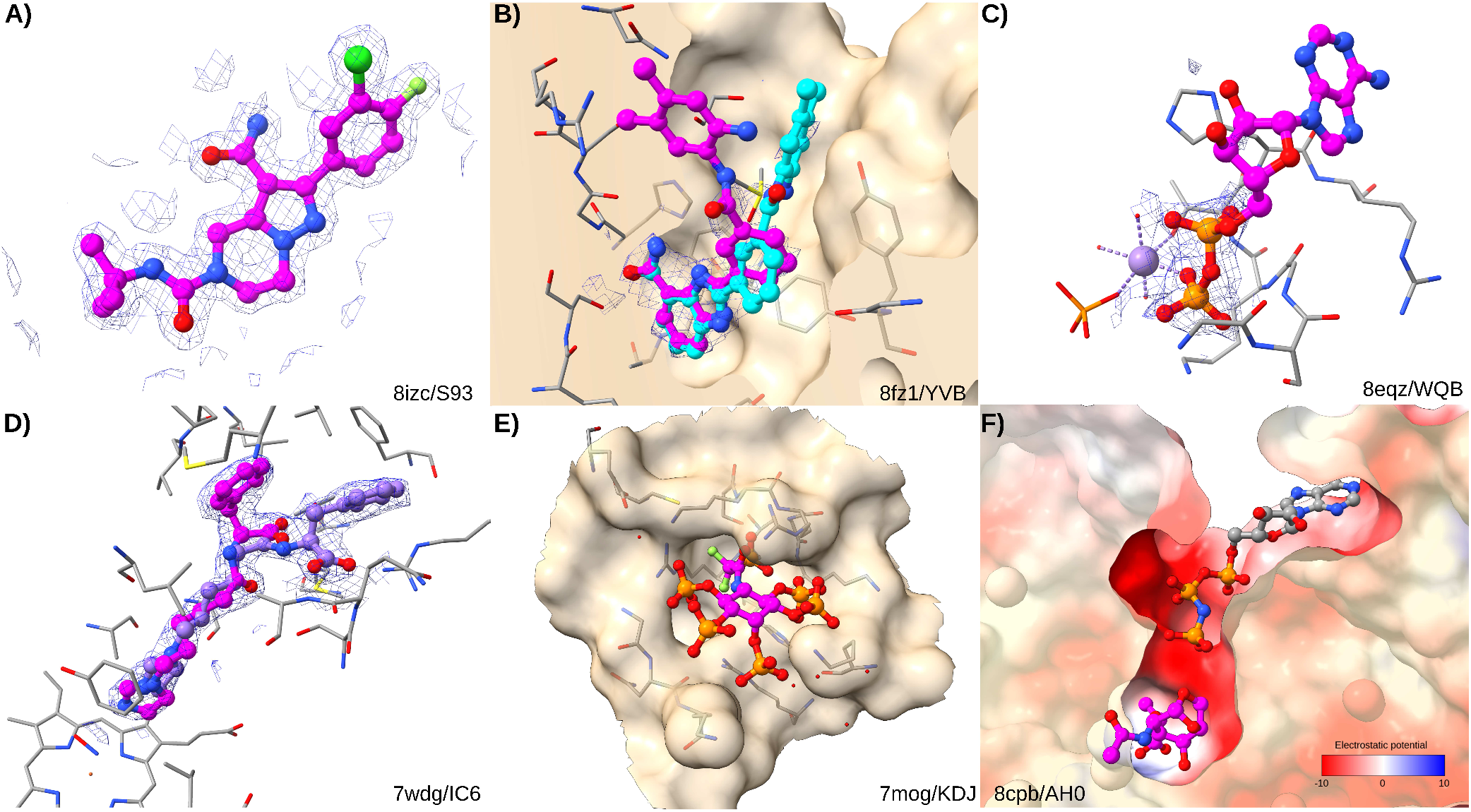
Example ligand–protein complexes from RNP set. The ligand of interest is highlighted by magenta carbon atoms. Electron density maps are shown at 1σ. A) Example of a well-defined ligand with an EDIA_m_ value of 1.10 (PDB ID 8icz/CCD S93). B) Example of a partially defined ligand (8fz1/YVB) with EDIA_m_ value of 0.09. The best docking pose of AC (cyan) is defined as failure but displays as good a density fit as the ligand conformation in the PDB model. C) Example of a partially defined ligand (8eqz/WQB) with EDIA_m_ value of 0.14. D) Example complex with two alternative ligand conformations, both present as distinct cases in the RNP set (7wdg/IC6). E) Example of a non-druglike highly phosphorylated ligand carrying a net charge of −12 (7mog/KDJ). F) Example of an unphysical PDB model (8cpb/AH0). The protein has a strongly negative electrostatic potential in the binding site and is additionally bound to a cofactor carrying a charge of −4. The negatively charged ligand cannot bind to this site in the absence of additional cations. All figures were generated with UCSF ChimeraX.^32,33^

We also calculated the optimal intuitively interpretable structural tool for evaluating real-world relevance (OiiSTER) map values^43^ and categories for all ligands (Fig. S3). OiiSTER assesses the molecular beauty of synthetic or virtual small molecules by their structural complexity and usuality, dividing complexity into minimal/regular/over-elaborated categories and usuality into trivial/balanced/unusual categories. This analysis shows that Astex has the highest amount of the regular/balanced class, which is the most relevant for drug design, while the RNP set has the highest amount of unusual ligands. The RNP set contains a high proportion of co-factors, often featuring phosphate groups, which are highly over-represented with respect to drug-like molecules, i.e. 672 out of 2,812 ligands (24%) contain at least one phosphorus atom (examples shown in Fig. 2C,E). For comparison, only 2 ligands of the Astex set (2%) and 12 of the PDBbind set (4%) contain phosphorus atoms. A significant part of these ligands are labeled as “prevalent” ligands by the RNP authors, defined as ligands with more than 100 ligands in the training set having a >0.9 RDKit topological fingerprint Tanimoto similarity.^57^ Their over-representation can therefore be reduced by filtering out prevalent ligands.

Furthermore, 26% of all RNP cases belong to the 10 most populated structural clusters alone, demonstrating an imbalance in the data set, which is absent in the Astex and PDBbind sets due to removal of similar complexes.

We further analyzed RNP ligand properties as a function of binding site and ligand pose similarities separately (Fig. 3). The analysis shows a strong data heterogeneity and hidden correlations in the dataset, complicating one-dimensional analyses. For example, ligands with a low ligand pose similarity are generally large, flexible, less well-defined, and less buried than ligands with a high ligand pose similarity. More than 80% of the complexes have a binding site similarity above 80%, and the ligands in these bins are relatively strongly buried and well-defined by the density.

**Figure 3.**
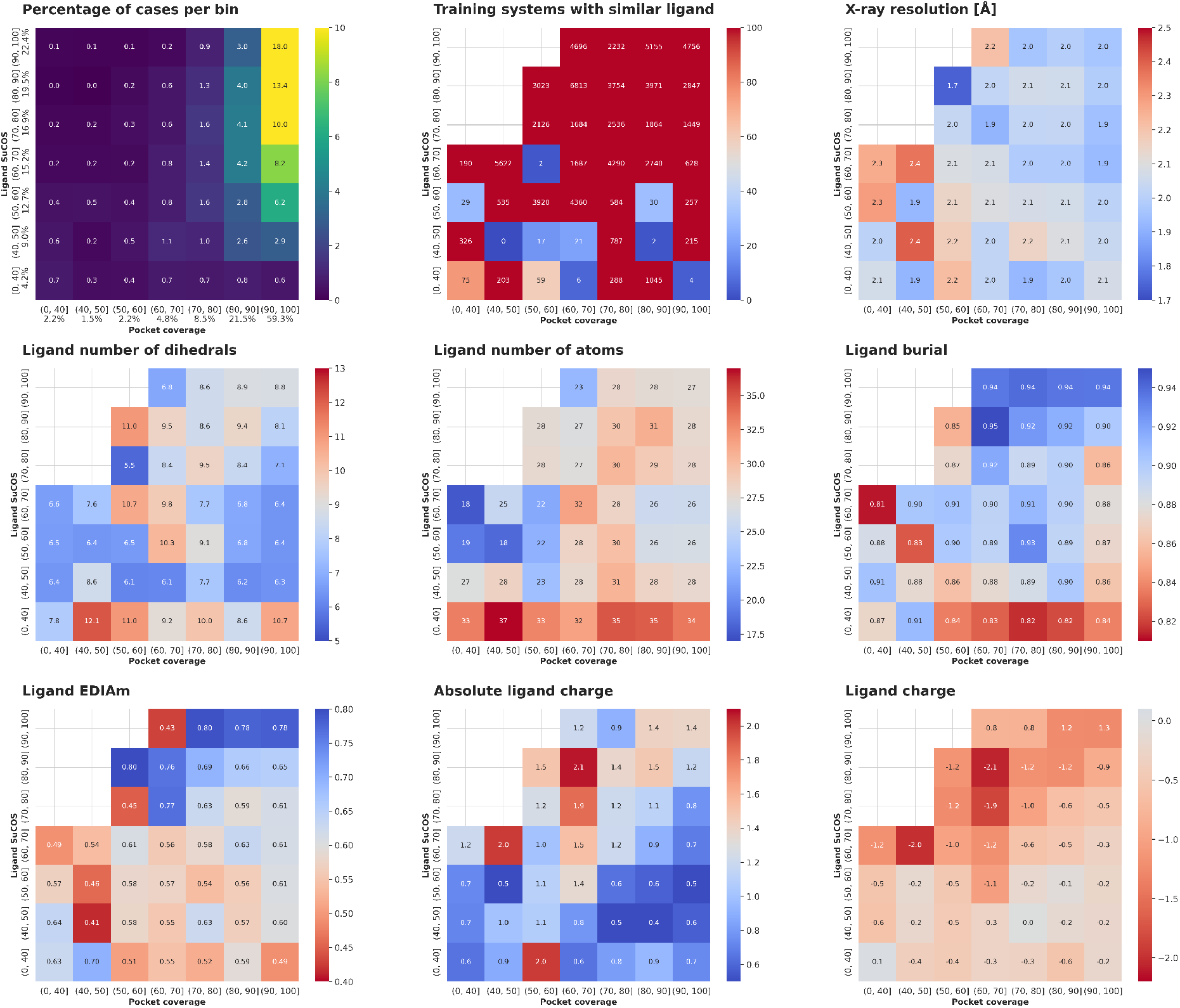
Properties of the RNP complexes as a function of binding site (pocket coverage) and ligand pose (SuCOS score) similarities. The first plot shows the percentage of cases in each bin. Bins with less than 5 cases are omitted for clarity in the other plots. The corresponding analysis for the RNP-F set can be found in Fig. S4, showing a more balanced distribution of properties.

To improve the quality, diversity, and balance of the RNP set, we applied the following filters in the given order:

- Density support of ligand (EDIA_m_) ≥ 0.4
- Density support of ligand environment within 5 Å ≥ 0.4
- Maximally 100 analogous ligands in the training set
- Maximally one complex per PDB ID, keeping the one with lowest similarity score
- Maximally 5 complexes belonging to the same cluster, keeping the ones with lowest similarity score

This results in the RNP-F set of 919 cases spread across 516 clusters. As shown in Figure 1, RNP-F removes a number of problematic cases but remains more challenging than both Astex and PDBbind sets in terms of ligand flexibility and solvent exposure. 9% of ligands in the RNP-F set contain at least one phosphorus atom, still more than in the Astex (2%) and PDBbind (4%) set, but significantly less than in the RNP (24%) set.

### 3.2 Evaluation of Docking Results

We carried out re-docking calculations on the RNP set using different algorithms and parameters. Figure 4A shows that the success rate at 2 Å RMSD of the highest ranked pose on the full RNP set is 54–59% for AC with different sampling parameters and 33–43% for AutoDock Vina and derivatives. These rates increase to 63–68% for AC and to 40-48% for Vina and derivatives when considering the filtered RNP-F set (Fig. 4B). These success rates are lower than the ones for the Astex and PDBbind set^17^ due to the higher ligand flexibility and lower ligand density support. Analyzing AC_norm_ and Vina with exhaustivity 100 in mode detail, the stratification (Fig. 4C–F) demonstrates that both docking approaches perform worse for very flexible ligands, highly solvent exposed ligands, poorly defined ligand poses, and ligands carrying more than one net charge. AC_norm_ systematically outperforms Vina in all categories, in-line with previous results.^17^

**Figure 4.**
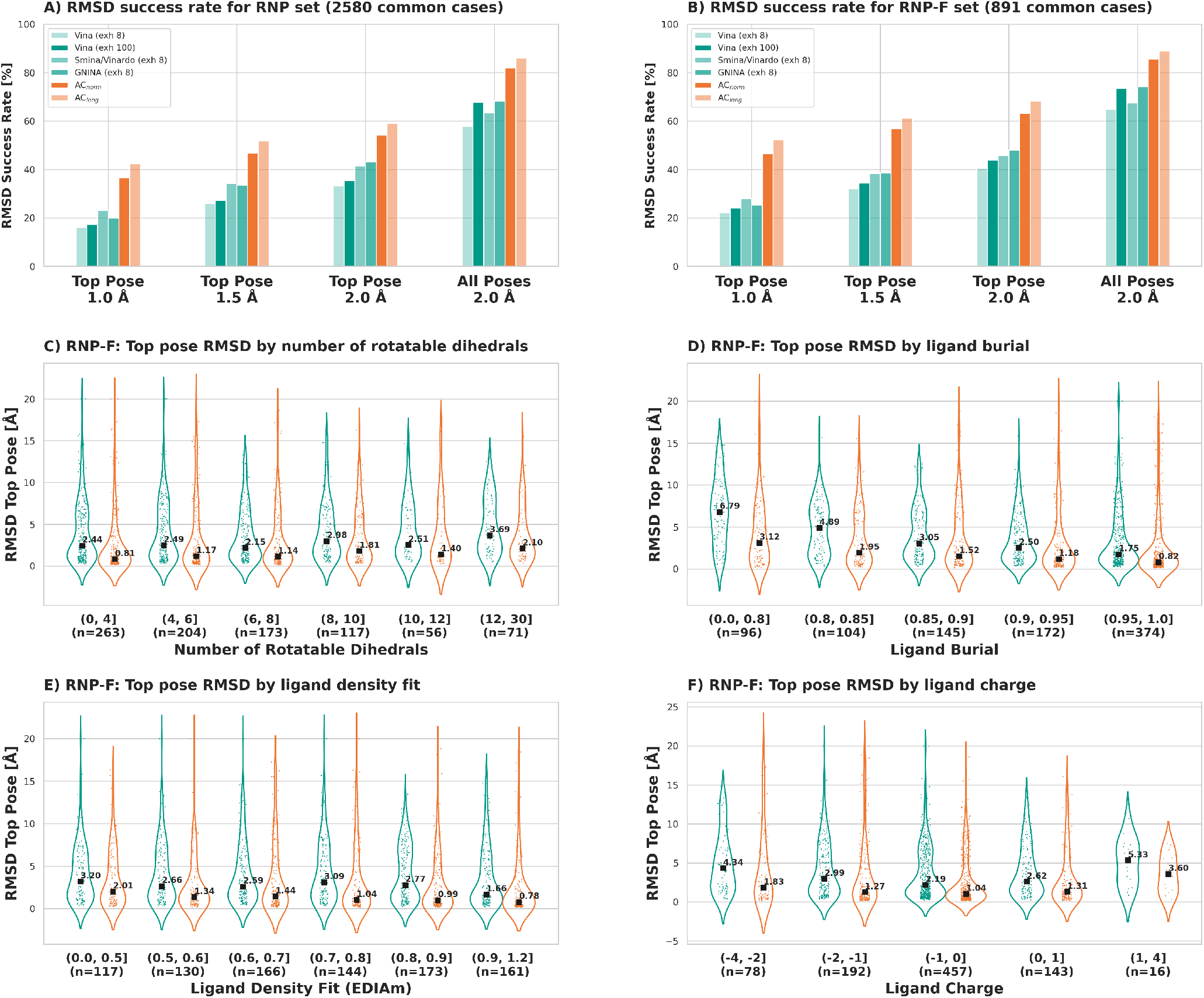
Analysis of re-docking results for AutoDock Vina and forks (green) and for AC (orange). A) RMSD success rates for RNP set common cases. B) RMSD success rates for RNP-F set common cases. Success rates at a RMSD cutoff of 1.0, 1.5, and 2.0 Å for the top pose and at 2.0 Å for all poses independent of their rank are given. For clarity, in part C to F only the results of AC_norm_ and Vina (exh 100) for the RNP-F set are shown. C) Top pose RMSD as a function of the number of rotatable dihedrals in the ligand. Median values are shown by black squares and annotated. D) Top pose RMSD as a function of ligand burial. E) Top pose RMSD as a function of ligand density fit (EDIA_m_). F) Top pose RMSD as a function of ligand charge.

We found that all methods yielded more scoring failures on the RNP set as compared to the Astex and PDBbind sets, and that the ratio was lower in the RNP-F set than in the RNP set (Tab. 1). In the past, we found that about one third of scoring failures of the AC scoring function could be attributed to issues in the scoring function (mainly for metal-bound and very exposed ligands), one third to issues in structure preparation (errors in chemical ligand structure, wrong protonation states or tautomers, missing cofactors), and one third to issues in the structual model in the PDB (missing electron density support of ligand, unassigned extra density, crystal contacts).^17^ The RNP set displayed the highest ratio of scoring failures among all sets for all four scoring functions, reaching even more than 50% for Vina. The filtered RNP-F set has less scoring failures, at a similar level as the PDBbind set. On average, complexes resulting in scoring failures with all scoring functions tend to have smaller and less flexible ligands, which are more solvent exposed, have a higher B-factor, are more charged, and more ill-defined by the electron density than consistent scoring successes (Fig. S5). An example of a scoring failure in all 4 scoring functions, which is obviously due to an issue in the structural model, is shown in Fig. 2F. Here, the protein binding site displays a strongly negative electrostatic potential, further enhanced by the cofactor carrying a net charge of −4, making it impossible for the negatively charged ligand to bind to this site in the absence of additional cations.

**Table 1.** Scoring failures (ScFail) of 4 scoring functions for different benchamrk sets. The intersection of scoring failures and scoring successes (ScSucc) are also given where calculated. ND: not determined.

|  | Astex [%] | PDBbind [%] | RNP [%] | RNP-F [%] |
| --- | --- | --- | --- | --- |
| ScFail AC | 10.6 | 24.2 | 33.4 | 26.0 |
| ScFail Vina | 30.6 | 40.4 | 51.6 | 42.0 |
| ScFail Vinardo | ND | ND | 33.1 | 29.2 |
| ScFail GNINA | ND | ND | 24.2 | 26.3 |
| ∩ ScFail | ND | ND | 7.2 | 6.2 |
| ∩ ScSucc | ND | ND | 30.7 | 38.4 |

### 3.3 Comparison of Docking and Co-Folding Results

When considering the full RNP dataset, AlphaFold 3 outperforms the physics-based docking tools for cases that are similar to cases from its training set, but Vina and especially AC perform better in the low-data regime (Fig. 5A). For the cleaned RNP-F set, these trends are even more evident (Fig. 5B). The small decrease of AC and Vina success rates for the second similarity bin (20-30) in Fig. 5A is due to the fact that complexes in this bin have ligands that are on average ill-defined by the electron density (low ligand EDIA_m_ value) and very solvent exposed (see Fig. S6). As the first are filtered out in the RNP-F set, the dip is smaller in Fig. 5B. The AlphaFold 3 success rate indeed depends both on the binding pocket and the ligand pose similarity to the training set, while this is not the case for AC and Vina (Fig. 5C). Instead, as shown before, AC and Vina success rates correlate with the ligand burial and flexibility (Fig. 5D).

**Figure 5.**
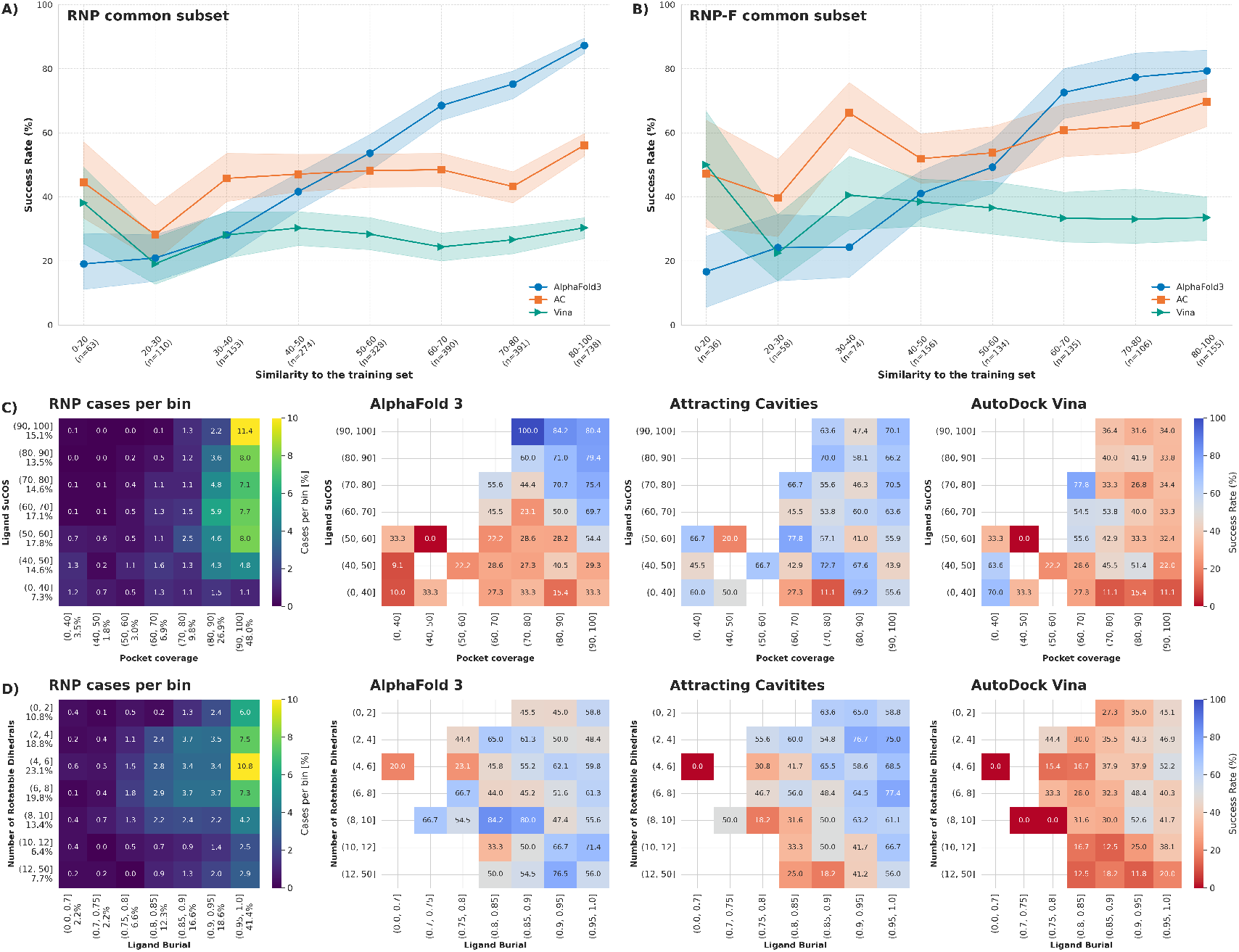
Comparison of success rates of AlphaFold 3, AC, and Vina. A) Success rates on RNP common subset (2,448 cases) plotted as in Ref.^10^. Shaded regions correspond to the 95% confidence interval, calculated from 1,000 bootstrap samples for each bin and method. B) Success rates on RNP-F common subset (854 cases). C) Success rates on RNP common subset as a function of binding site and ligand pose similarity separately. The plot on the left shows the percentage of cases in each bin. D) Success rates on RNP common subset as a function of ligand burial and number of rotatable dihedrals. For the RNP-F set, analogous graphs to parts C and D are shown in Fig. S7.

The same picture emerges from descriptive logistic regression models, calculating successful structure predictions as a function of two similarity and three physical variables after normalization to the range [0, 1]. (Tab. 2 and Tab. S1–S3). While AlphaFold 3 shows the largest coefficients and coefficient/error ratios for the two similarity variables, the opposite is true for AC and Vina. The large coefficient of the AlphaFold 3 model for the ligand density fit is due to the positive correlation of this variable to complex similarity (Fig. 3 and Fig. S6).

**Table 2.** Logistic regression model coefficients (Coef.) and standard errors (SE) for AlphaFold 3, Attracting Cavities, and AutoDock Vina success on the RNP set (2,448 common cases). All variables were rescaled to the range [0, 1] before regression. The accuracy and area under the ROC curve (AUC) for each model are given. Coefficients with a P-value <0.001 are marked in bold.

|  | AF3 |  | AC |  | Vina |  |
| --- | --- | --- | --- | --- | --- | --- |
|  | Coef. | SE | Coef. | SE | Coef. | SE |
| Ligand Pose Similarity | **4.106** | 0.29 | 0.327 | 0.25 | 0.056 | 0.28 |
| Binding Pocket Similarity | **2.210** | 0.35 | -0.004 | 0.30 | -0.859 | 0.32 |
| Ligand Burial | 0.548 | 0.29 | **3.457** | 0.29 | **3.656** | 0.35 |
| Ligand Flexibility | 0.753 | 0.28 | **-1.435** | 0.23 | **-1.045** | 0.25 |
| Ligand Density Fit | **2.032** | 0.21 | **1.444** | 0.20 | **1.053** | 0.22 |
| Constant | **-5.947** | 0.42 | **-3.446** | 0.35 | **-3.541** | 0.39 |
| Accuracy | 0.744 |  | 0.637 |  | 0.668 |  |
| AUC | 0.79 |  | 0.70 |  | 0.67 |  |

## 4 Summary and Conclusions

The aim of this work was to comparatively assess the usefulness of co-folding and docking approaches for small-molecule drug design. These two approaches are fundamentally different and therefore difficult to compare. However, a comparison is warranted because both are used for ligand binding pose prediction in structure-based drug design, and a user should be able to make an informed choice of method. To this end, we used the recently published and highly annotated Runs N’ Poses set of ligand–protein complexes, with which it was shown that current co-folding methods work well for predicting complexes similar to the training data but fail to generalize to novel data.^10^ No comparison to physics-based docking methods was provided by the authors.

Our analysis of the RNP set, which has been developed to detect data leakage in co-folding methods, showed that it should be further optimized for benchmarking pose prediction algorithms, as it contains (1) many ligands and binding pockets ill-defined by the electron density, (2) a very high percentage of non-druglike ligands, and (3) many complexes belonging to just a few clusters of similar structures. We therefore filtered this set to generate a more balanced and better quality reference set, RNP-F, composed of 919 complexes spread across 516 structural clusters. Filtering also reduced the high prevalence of phosphorus-containing ligands (from 24% in RNP to 9% in RNP-F), which are rare among small-molecule drugs.^58^

We carried out docking calculations on the RNP set with AutoDock Vina and its Smina/Vinardo and GNINA forks as well as our Attracting Cavities algorithm. As we do local re-docking on predefined binding pockets, success rates should be considered an upper bound to what can be expected in cross-docking. In this context it should be remembered that physics-based docking algorithms have proven their merit in countless successful drug design projects over the last decades. Their assessment on known ligand–protein complexes from the PDB has always been just a necessary first step in their development, but their true value is shown by their predictive capabilities in real-world utilization.

The docking results demonstrate that AC outperforms all Vina-based algorithms albeit at longer CPU times, and that the quality of docking predictions depends on ligand burial, ligand flexibility, and ligand density fit. One fourth to one half of docking failures are not due to insufficient sampling but to the scoring functions ranking an alternative ligand binding pose better than the one described in the PDB model. Underlying reasons for scoring failures can be issues in the scoring function itself, issues in structure preparation, or issues in the PDB model. We give examples showing how docking results yield insight into the validity of structural data in the PDB, which are error-prone models fitted to experimental data and which should not be considered as “ground truth.”

Finally, we compare AlphaFold 3 predictions with docking results from AC and Vina, demonstrating that physics-based docking outperforms co-folding for novel complexes with low similarity to complexes in the PDB, which is obviously the predominant application domain for a ligand binding pose prediction algorithm in a drug design context, aimed at discovering new and diverse ligands for underexplored targets. Docking success rates are determined by physical ligand properties such as flexibility and binding site burial, while co-folding success rates are mainly determined by binding site and ligand pose similarity to the training set. The big advantage of co-folding methods over docking methods is that the first need only the protein sequence and chemical ligand structure as input, while the latter necessitate a 3D target structure and, at least for practical purposes, information on the localization of the binding site. When this information is not readily available, a combination of co-folding and docking methods may be the most promising approach for drug design.

In summary, ligand binding pose prediction for small-molecule drug design, which is concerned with novel ligands and novel binding pockets, is a problem not yet solved by co-folding methods and better tackled with physics-based methods.

## 5 Data and Software Availability

Software and procedures mentioned can be accessed on the following websites: Docking input files, results, and annotations from the present work (https://doi.org/10.5281/zenodo.17865538), Runs N’ Poses data set (https://github.com/plinder-org/runs-n-poses, https://zenodo.org/records/16754298), MGL tools for generating Vina input files (https://ccsb.scripps.edu/mgltools/), AutoDock Vina docking program (https://vina.scripps.edu), Smina docking program (https://github.com/mwojcikowski/smina), GNINA docking program (https://github.com/gnina/gnina), CHARMM molecular simulation program (https://www.charmm.org), density-fitness to calculate EDIA_m_ values (https://github.com/PDB-REDO/density-fitness), sPyRMSD to calculate symmetry-adapted RMSD values (https://github.com/RMeli/spyrmsd), RDKit (https://www.rdkit.org/), Open Babel version 2.4.1 for ligand coordinate manipulation (https://openbabel.org), statsmodel python module for logistic regression (https://www.statsmodels.org), scikit-learn python module for logisitic regression (https://scikit-learn.org), SwissParam server for generating ligand force fields for AC (https://www.swissparam.ch/), UCSF ChimeraX software for molecular analysis and visualization (https://www.cgl.ucsf.edu/chimerax).

## References

1. Abramson, J. et al. Accurate structure prediction of biomolecular interactions with AlphaFold 3. Nature 630, 493–500, DOI: 10.1038/s41586-024-07487-w (2024).

2. Bryant, P., Kelkar, A., Guljas, A., Clementi, C. & Noé, F. Structure prediction of protein-ligand complexes from sequence information with Umol. Nat. Commun. 15, 4536, DOI: 10.1038/s41467-024-48837-6 (2024).

3. Krishna, R. et al. Generalized biomolecular modeling and design with RoseTTAFold All-Atom. Science 384, eadl2528, DOI: 10.1126/science.adl2528 (2024).

4. Chai Discovery. Chai-1: Decoding the molecular interactions of life. bioRxiv DOI: 10.1101/2024.10.10.615955 (2024).

5. Wohlwend, J. et al. Boltz-1: Democratizing biomolecular interaction modeling. bioRxiv DOI: 10.1101/2024.11.19.624167 (2024).

6. Passaro, S. et al. Boltz-2: Towards accurate and efficient binding affinity prediction. bioRxiv DOI: 10.1101/2025.06.14.659707 (2025).

7. Stark, H. et al. BoltzGen: Toward Universal Binder Design. bioRxiv 2025.11.20.689494, DOI: 10.1101/2025.11.20.689494 (2025).

8. ByteDance AML AI4Science Team et al. Protenix – advancing structure prediction through a comprehensive AlphaFold3 reproduction. bioRxiv DOI: 10.1101/2025.01.08.631967 (2025).

9. Qiao, Z., Nie, W., Vahdat, A., Miller, T. F. & Anandkumar, A. State-specific protein–ligand complex structure prediction with a multiscale deep generative model. Nat. Mach. Intell. 6, 195–208, DOI: 10.1038/s42256-024-00792-z (2024).

10. Škrinjar, P., Eberhardt, J., Tauriello, G., Schwede, T. & Durairaj, J. Have protein-ligand cofolding methods moved beyond memorisation? bioRxiv 2025.02.03.636309, DOI: 10.1101/2025.02.03.636309 (2025).

11. Masters, M. R., Mahmoud, A. H. & Lill, M. A. Investigating whether deep learning models for co-folding learn the physics of protein-ligand interactions. Nat. Commun. 16, 8854, DOI: 10.1038/s41467-025-63947-5 (2025).

12. Paggi, J. M., Pandit, A. & Dror, R. O. The Art and Science of Molecular Docking. Annu. Rev. Biochem. 93, 389–410, DOI: 10.1146/annurev-biochem-030222-120000 (2024).

13. Corso, G., Stärk, H., Jing, B., Barzilay, R. & Jaakkola, T. DiffDock: Diffusion Steps, Twists, and Turns for Molecular Docking. arXiv 2210.01776, DOI: 10.48550/arXiv.2210.01776 (2023).

14. Hartshorn, M. J. et al. Diverse, high-quality test set for the validation of protein-ligand docking performance. J. Med. Chem. 50, 726–741, DOI: 10.1021/jm061277y (2007).

15. Su, M. et al. Comparative Assessment of Scoring Functions: The CASF-2016 Update. J. Chem. Inf. Model. 59, 895–913, DOI: 10.1021/acs.jcim.8b00545 (2019).

16. Zoete, V. et al. Attracting cavities for docking. Replacing the rough energy landscape of the protein by a smooth attracting landscape. J. Comp. Chem. 37, 437–447, DOI: 10.1002/jcc.24249 (2016).

17. Röhrig, U. F., Goullieux, M., Bugnon, M. & Zoete, V. Attracting cavities 2.0: Improving the flexibility and robustness for small-molecule docking. J. Chem. Inf. Model. 63, 3925–3940, DOI: 10.1021/acs.jcim.3c00054 (2023).

18. Goullieux, M., Zoete, V. & Röhrig, U. F. Two-step covalent docking with attracting cavities. J. Chem. Inf. Model. 63, 7847–7859 (2023).

19. Trott, O. & Olson, A. J. AutoDock Vina: improving the speed and accuracy of docking with a new scoring function, efficient optimization, and multithreading. J. Comput. Chem. 31, 455–461, DOI: 10.1002/jcc.21334 (2010).

20. Eberhardt, J., Santos-Martins, D., Tillack, A. F. & Forli, S. AutoDock Vina 1.2.0: New Docking Methods, Expanded Force Field, and Python Bindings. J. Chem. Inf. Model. 61, 3891–3898, DOI: 10.1021/acs.jcim.1c00203 (2021).

21. Brooks, B. R. et al. CHARMM: the biomolecular simulation program. J. Comput. Chem. 30, 1545–1614, DOI: 10.1002/jcc.21287 (2009).

22. Huang, J. et al. CHARMM36m: an improved force field for folded and intrinsically disordered proteins. Nat Methods 14, 71–73, DOI: 10.1038/nmeth.4067 (2017).

23. Haberthür, U. & Caflisch, A. Facts: Fast analytical continuum treatment of solvation. J. Comput. Chem. 29, 701–715, DOI: 10.1002/jcc.20832 (2008).

24. Bugnon, M. et al. SwissParam 2023: A Modern Web-Based Tool for Efficient Small Molecule Parametrization. J. Chem. Inf. Model. 63, 6469–6475, DOI: 10.1021/acs.jcim.3c01053 (2023).

25. Koes, D. R., Baumgartner, M. P. & Camacho, C. J. Lessons Learned in Empirical Scoring with smina from the CSAR 2011 Benchmarking Exercise. J. Chem. Inf. Model. 53, 1893–1904, DOI: 10.1021/ci300604z (2013).

26. Quiroga, R. & Villarreal, M. A. Vinardo: A Scoring Function Based on Autodock Vina Improves Scoring, Docking, and Virtual Screening. PLoS One 11, e0155183, DOI: 10.1371/journal.pone.0155183 (2016). 27171006.

27. McNutt, A. T., Li, Y., Meli, R., Aggarwal, R. & Koes, D. R. GNINA 1.3: The next increment in molecular docking with deep learning. J Cheminf 17, 1–8, DOI: 10.1186/s13321-025-00973-x (2025).

28. wwPDB consortium. Protein Data Bank: the single global archive for 3D macromolecular structure data. Nucleic Acids Res. 47, D520–D528, DOI: 10.1093/nar/gky949 (2019).

29. Burley, S. K. et al. Protein Data Bank (PDB): The Single Global Macromolecular Structure Archive. In Wlodawer, A., Dauter, Z. & Jaskolski, M. (eds.) Protein Crystallography: Methods and Protocols, 627–641, DOI: 10.1007/978-1-4939-7000-1_26 (Springer, New York, NY, 2017).

30. Durairaj, J. et al. PLINDER: The protein-ligand interactions dataset and evaluation resource. bioRxiv 2024.07.17.603955, DOI: 10.1101/2024.07.17.603955 (2024).

31. Leung, S., Bodkin, M., von Delft, F., Brennan, P. & Morris, G. SuCOS is Better than RMSD for Evaluating Fragment Elaboration and Docking Poses. ChemRxiv DOI: 10.26434/chemrxiv.8100203.v1 (2019).

32. Pettersen, E. F. et al. UCSF ChimeraX: Structure visualization for researchers, educators, and developers. Protein Sci. 30, 70–82, DOI: 10.1002/pro.3943 (2021).

33. Meng, E. C. et al. UCSF ChimeraX: Tools for structure building and analysis. Protein Sci. 32, e4792, DOI: 10.1002/pro.4792 (2023).

34. van Ginkel, G. et al. PDBeCIF: An open-source mmCIF/CIF parsing and processing package. BMC Bioinf 22, 1–7, DOI: 10.1186/s12859-021-04271-9 (2021).

35. RDKit: Open-source cheminformatics software. http://www.rdkit.org/.

36. O’Boyle, N. M. et al. Open babel: An open chemical toolbox. J. Cheminf. 3, 33, DOI: 10.1186/1758-2946-3-33 (2011).

37. Morris, G. M. et al. AutoDock4 and AutoDockTools4: automated docking with selective receptor flexiblity. J. Comput. Chem. 30, 2785–91, DOI: 10.1002/jcc.21256 (2009).

38. Bugnon, M. et al. SwissDock 2024: Major enhancements for small-molecule docking with Attracting Cavities and AutoDock Vina. Nucleic Acids Res. 52, W324–W332, DOI: 10.1093/nar/gkae300 (2024).

39. Best, R. B. et al. Optimization of the additive CHARMM all-atom protein force field targeting improved sampling of the backbone φ, ψ and side-chain χ1 and χ2 dihedral angles. J. Chem. Theory Comput. 8, 3257–3273, DOI: 10.1021/ct300400x (2012).

40. Meyder, A., Nittinger, E., Lange, G., Klein, R. & Rarey, M. Estimating Electron Density Support for Individual Atoms and Molecular Fragments in X-ray Structures. J. Chem. Inf. Model. 57, 2437–2447, DOI: 10.1021/acs.jcim.7b00391 (2017).

41. density-fitness, part of the PDB-REDO tools for macromolecular structure optimisation and analysis, https://github.com/PDB-REDO/density-fitness.

42. Joosten, R. P., Long, F., Murshudov, G. N. & Perrakis, A. The PDB_REDO server for macromolecular structure model optimization. IUCrJ 1, 213–220, DOI: 10.1107/S2052252514009324 (2014).

43. Daina, A. & Zoete, V. Rethinking molecular beauty in the deep learning era. bioRxiv DOI: 10.64898/2025.12.03.692079 (2025).

44. Meli, R. & Biggin, P. C. Spyrmsd: Symmetry-corrected RMSD calculations in Python. J Cheminf 12, 49, DOI: 10.1186/s13321-020-00455-2 (2020-12).

45. Robin, X. et al. Assessment of protein–ligand complexes in CASP15. Proteins: Struct, Funct, Bioinf 91, 1811–1821, DOI: 10.1002/prot.26601 (2023).

46. Biasini, M. et al. OpenStructure: An integrated software framework for computational structural biology. Acta Crystallogr. Sect. D: Biol. Crystallogr. 69, 701–709, DOI: 10.1107/S0907444913007051 (2013).

47. Pedregosa, F. et al. Scikit-learn: Machine learning in Python. J. Mach. Learn. Res. 12, 2825–2830 (2011).

48. Seabold, S. & Perktold, J. statsmodels: Econometric and statistical modeling with python. In 9th Python in Science Conference (2010).

49. Neves, M. a. C., Totrov, M. & Abagyan, R. Docking and scoring with ICM: The benchmarking results and strategies for improvement. J. Comput. Aided. Mol. Des. 26, 675–86, DOI: 10.1007/s10822-012-9547-0 (2012).

50. Corbeil, C. R., Williams, C. I. & Labute, P. Variability in docking success rates due to dataset preparation. J. Comput. Aided. Mol. Des. 26, 775–86, DOI: 10.1007/s10822-012-9570-1 (2012).

51. Brozell, S. R. et al. Evaluation of DOCK 6 as a pose generation and database enrichment tool. J. Comput. Aided. Mol. Des. 26, 749–73, DOI: 10.1007/s10822-012-9565-y (2012). 22569593.

52. Repasky, M. P. et al. Docking performance of the glide program as evaluated on the Astex and DUD datasets: A complete set of glide SP results and selected results for a new scoring function integrating WaterMap and glide. J. Comput. Aided. Mol. Des. 26, 787–799, DOI: 10.1007/s10822-012-9575-9 (2012).

53. Santos-Martins, D. et al. Accelerating AutoDock4 with GPUs and Gradient-Based Local Search. J. Chem. Theory Comput. 17, 1060–1073, DOI: 10.1021/acs.jctc.0c01006 (2021).

54. Park, H., Zhou, G., Baek, M., Baker, D. & DiMaio, F. Force Field Optimization Guided by Small Molecule Crystal Lattice Data Enables Consistent Sub-Angstrom Protein–Ligand Docking. J. Chem. Theory Comput. 17, 2000–2010, DOI: 10.1021/acs.jctc.0c01184 (2021).

55. Liu, Z. et al. Forging the Basis for Developing Protein–Ligand Interaction Scoring Functions. Acc. Chem. Res. 50, 302–309, DOI: 10.1021/acs.accounts.6b00491 (2017).

56. Roehrig, U. F., Goullieux, M., Bugnon, M. & Zoete, V. Benchmark data for attracting cavities 2.0 small-molecular docking program. Zenodo DOI: 10.5281/zenodo.7940100 (2023).

57. Rogers, D. & Hahn, M. Extended-Connectivity Fingerprints. J. Chem. Inf. Model. 50, 742–754, DOI: 10.1021/ci100050t (2010).

58. Benedetto Tiz, D., Rosati, O. & Sancineto, L. Sulfur- and phosphorus-containing FDA approved drugs in the last five years (2020–2024): A journey among small molecules. Phosphorus, Sulfur, Silicon Relat. Elem. 1–39, DOI: 10.1080/10426507.2025.2496521 (2025).

