## Supplementary Data for "Comparative Assessment of the Utility of Co-Folding and Docking for Small-Molecule Drug Design"

### Supporting Tables

| | | | |
|---|---|---|---|
| **Dep. Variable:** | success_af3 | **No. Observations:** | 2446 |
| **Model:** | Logit | **Df Residuals:** | 2440 |
| **Method:** | MLE | **Df Model:** | 5 |
| **Date:** | Mon, 01 Dec 2025 | **Pseudo R-squ.:** | 0.2030 |
| **Time:** | 13:12:22 | **Log-Likelihood:** | -1270.8 |
| **converged:** | True | **LL-Null:** | -1594.4 |
| **Covariance Type:** | nonrobust | **LLR p-value:** | 1.169e-137 |

| | **coef** | **std err** | **z** | **P> \|z\|** | **[0.025** | **0.975]** |
|---|---|---|---|---|---|---|
| **Constant** | -5.9472 | 0.415 | -14.340 | 0.000 | -6.760 | -5.134 |
| **Ligand Burial** | 0.5480 | 0.292 | 1.879 | 0.060 | -0.024 | 1.120 |
| **Lig. No. Dihe.** | 0.7529 | 0.282 | 2.674 | 0.007 | 0.201 | 1.305 |
| **Ligand $EDIA_m$** | 2.0320 | 0.212 | 9.585 | 0.000 | 1.617 | 2.448 |
| **Pocket coverage** | 2.2099 | 0.351 | 6.291 | 0.000 | 1.521 | 2.898 |
| **Ligand SuCOS** | 4.1057 | 0.285 | 14.402 | 0.000 | 3.547 | 4.664 |

**Table S1.** Logistic Regression Results for AF3 Success. Accuracy 0.744, AUC 0.79.

| | | | |
|---|---|---|---|
| **Dep. Variable:** | success_ac | **No. Observations:** | 2446 |
| **Model:** | Logit | **Df Residuals:** | 2440 |
| **Method:** | MLE | **Df Model:** | 5 |
| **Date:** | Mon, 01 Dec 2025 | **Pseudo R-squ.:** | 0.09490 |
| **Time:** | 13:22:34 | **Log-Likelihood:** | -1533.6 |
| **converged:** | True | **LL-Null:** | -1694.4 |
| **Covariance Type:** | nonrobust | **LLR p-value:** | 2.262e-67 |

| | **coef** | **std err** | **z** | **P> \|z\|** | **[0.025** | **0.975]** |
|---|---|---|---|---|---|---|
| **Constant** | -3.4460 | 0.345 | -9.993 | 0.000 | -4.122 | -2.770 |
| **Ligand Burial** | 3.4574 | 0.290 | 11.912 | 0.000 | 2.888 | 4.026 |
| **Lig. No. Dihe.** | -1.4349 | 0.233 | -6.161 | 0.000 | -1.891 | -0.978 |
| **Ligand $EDIA_m$** | 1.4439 | 0.195 | 7.411 | 0.000 | 1.062 | 1.826 |
| **Pocket coverage** | -0.0036 | 0.301 | -0.012 | 0.991 | -0.593 | 0.586 |
| **Ligand SuCOS** | 0.3267 | 0.253 | 1.292 | 0.196 | -0.169 | 0.822 |

**Table S2.** Logistic Regression Results for AC Success ($AC_{norm}$) Accuracy 0.637, AUC 0.70.

| | | | |
|---|---|---|---|
| **Dep. Variable:** | success_vina | **No. Observations:** | 2446 |
| **Model:** | Logit | **Df Residuals:** | 2440 |
| **Method:** | MLE | **Df Model:** | 5 |
| **Date:** | Thu, 04 Dec 2025 | **Pseudo R-squ.:** | 0.07370 |
| **Time:** | 11:13:30 | **Log-Likelihood:** | -1345.3 |
| **converged:** | True | **LL-Null:** | -1452.4 |
| **Covariance Type:** | nonrobust | **LLR p-value:** | 2.744e-44 |

| | **coef** | **std err** | **z** | **P> \|z\|** | **[0.025** | **0.975]** |
|---|---|---|---|---|---|---|
| **Constant** | -3.5411 | 0.387 | -9.160 | 0.000 | -4.299 | -2.783 |
| **Ligand Burial** | 3.6596 | 0.346 | 10.569 | 0.000 | 2.981 | 4.338 |
| **Lig. No. Dihe.** | -1.0445 | 0.252 | -4.151 | 0.000 | -1.538 | -0.551 |
| **Ligand $\mathbf{EDIA_m}$** | 1.0533 | 0.218 | 4.830 | 0.000 | 0.626 | 1.481 |
| **Pocket coverage** | -0.8588 | 0.316 | -2.714 | 0.007 | -1.479 | -0.239 |
| **Ligand SuCOS** | 0.0564 | 0.276 | 0.204 | 0.838 | -0.485 | 0.598 |

**Table S3.** Logistic Regression Results for Vina Success. Accuracy 0.722, AUC 0.68.

### Supporting Figures

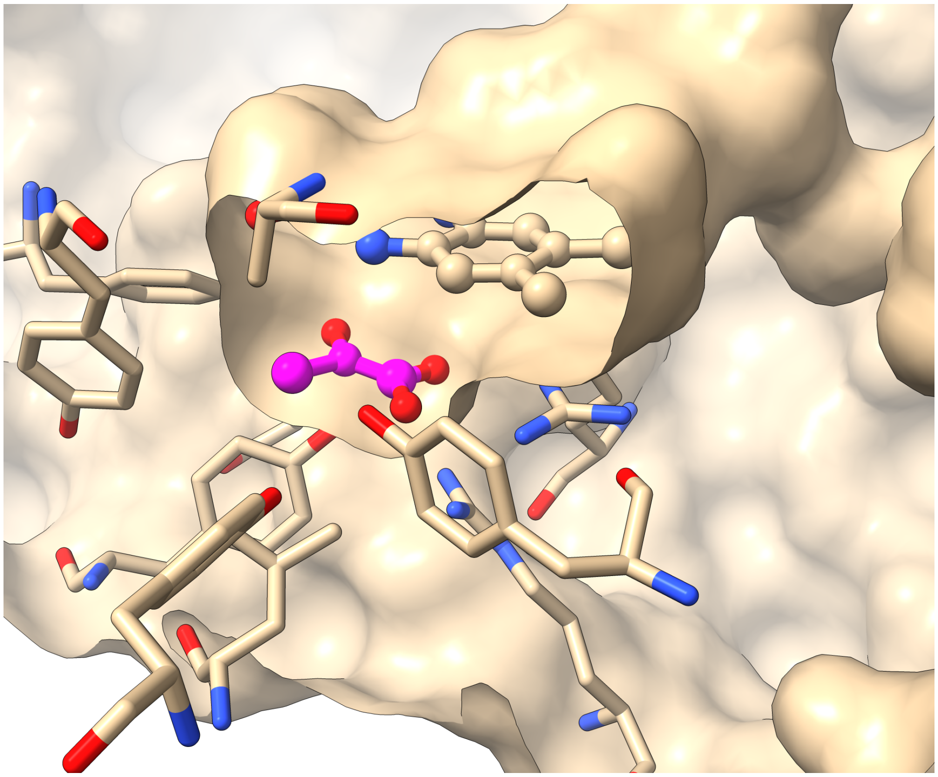

**Figure S1.** Example of Ligand Burial Calculations (PDB ID 7f22). For the pyruvate ligand (shown in magenta), which is included in the RNP set, ChimeraX gives a burial value of 0.55, CHARMM of 1.0, which corresponds better to this completely buried ligand pose.

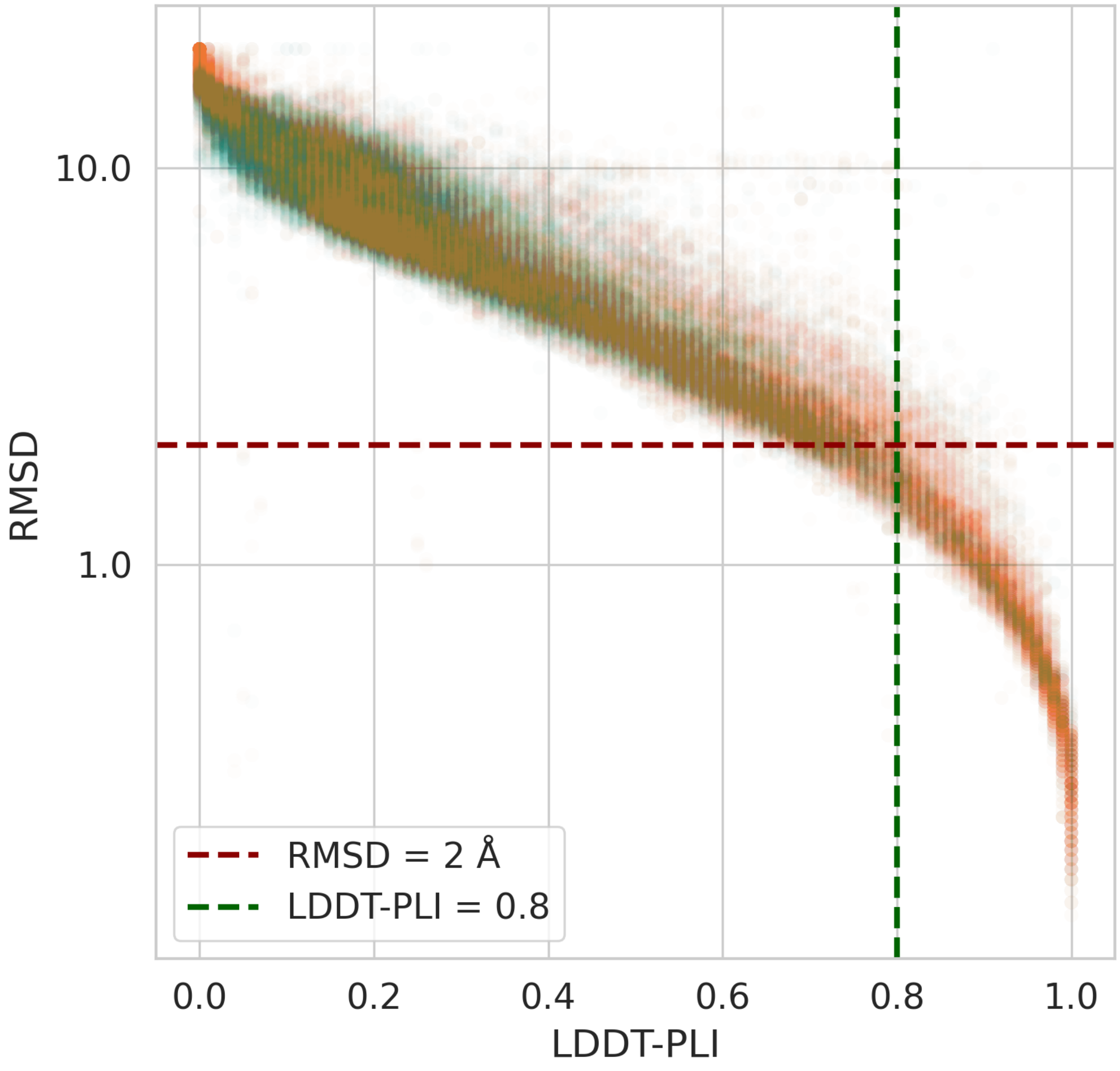

**Figure S2.** LDDT-PLI and RMSD values across all $AC_{norm}$ (orange) and Vina (exh. 100, green) docking predictions. Very few outliers with a low RMSD value but a low LDDT-PLI value are observed, compared to the same plot for co-folding results, Figure 7 from Ref.[1].

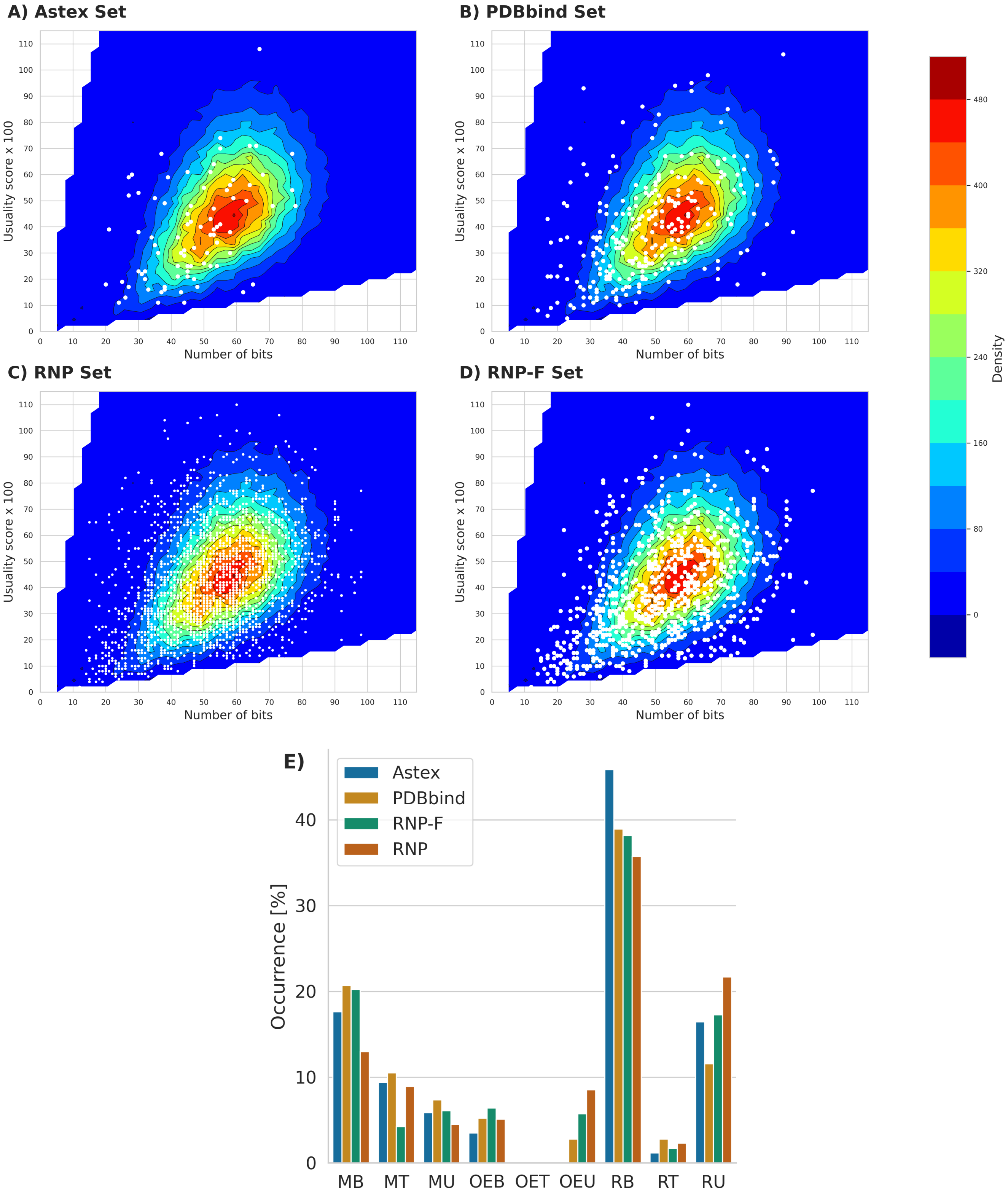

**Figure S3.** Oiister maps and categories[2] of ligand across benchmark sets. A) Astex set. B) PDBbind set. C) RNP set. D) RNP-F set. E) Oiister categories of all sets. Abbreviations: Complexity – minimal (M), regular (R), over-elaborated(OE); usuality – trivial (T), balanced (B), unusual (U).

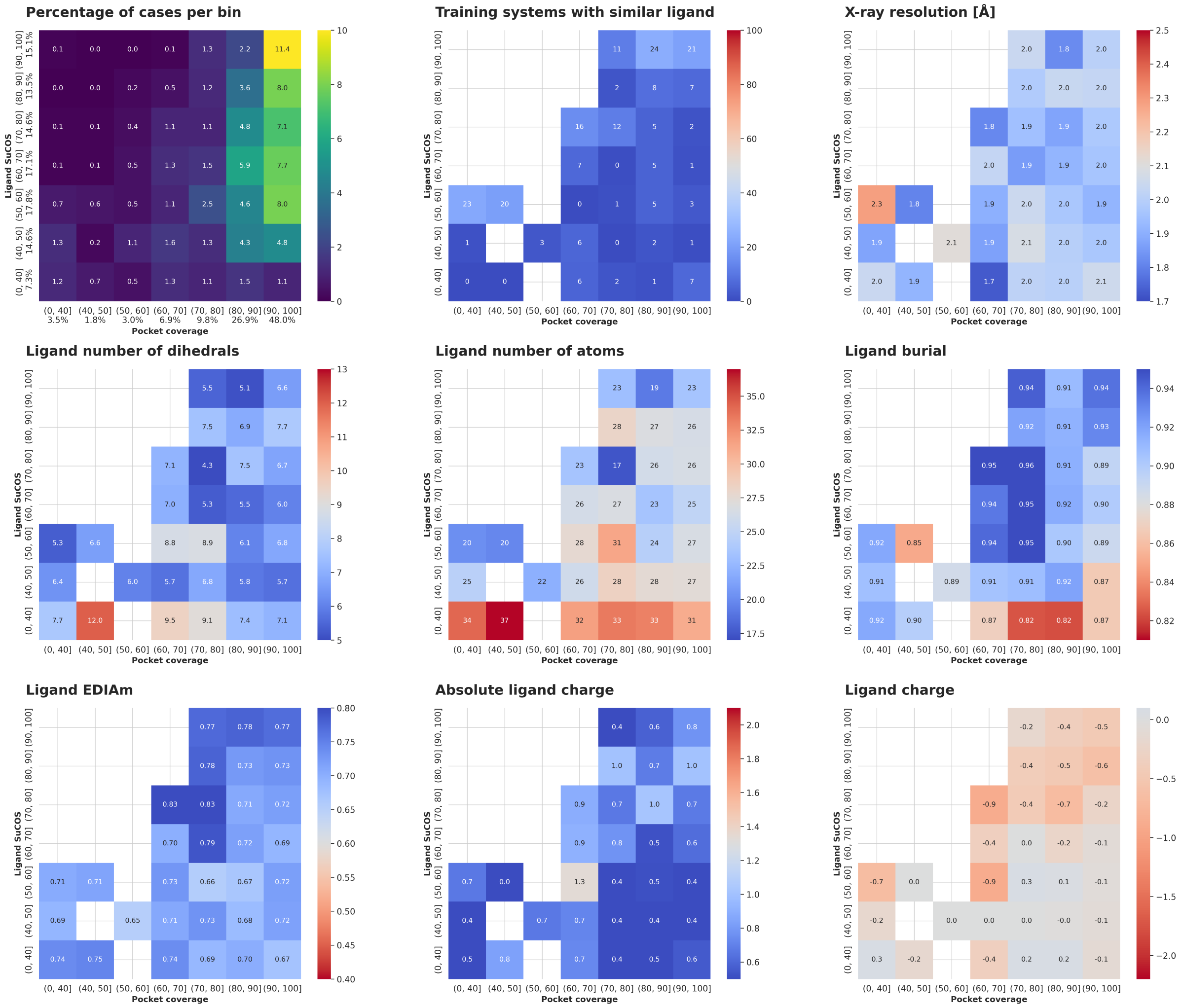

**Figure S4.** Properties of the RNP-F complexes as a function of binding site (pocket coverage) and ligand pose (SuCOS score) similarities. The first plot shows the percentage of cases in each bin. Bins with less than 5 cases are omitted for clarity in the other plots.

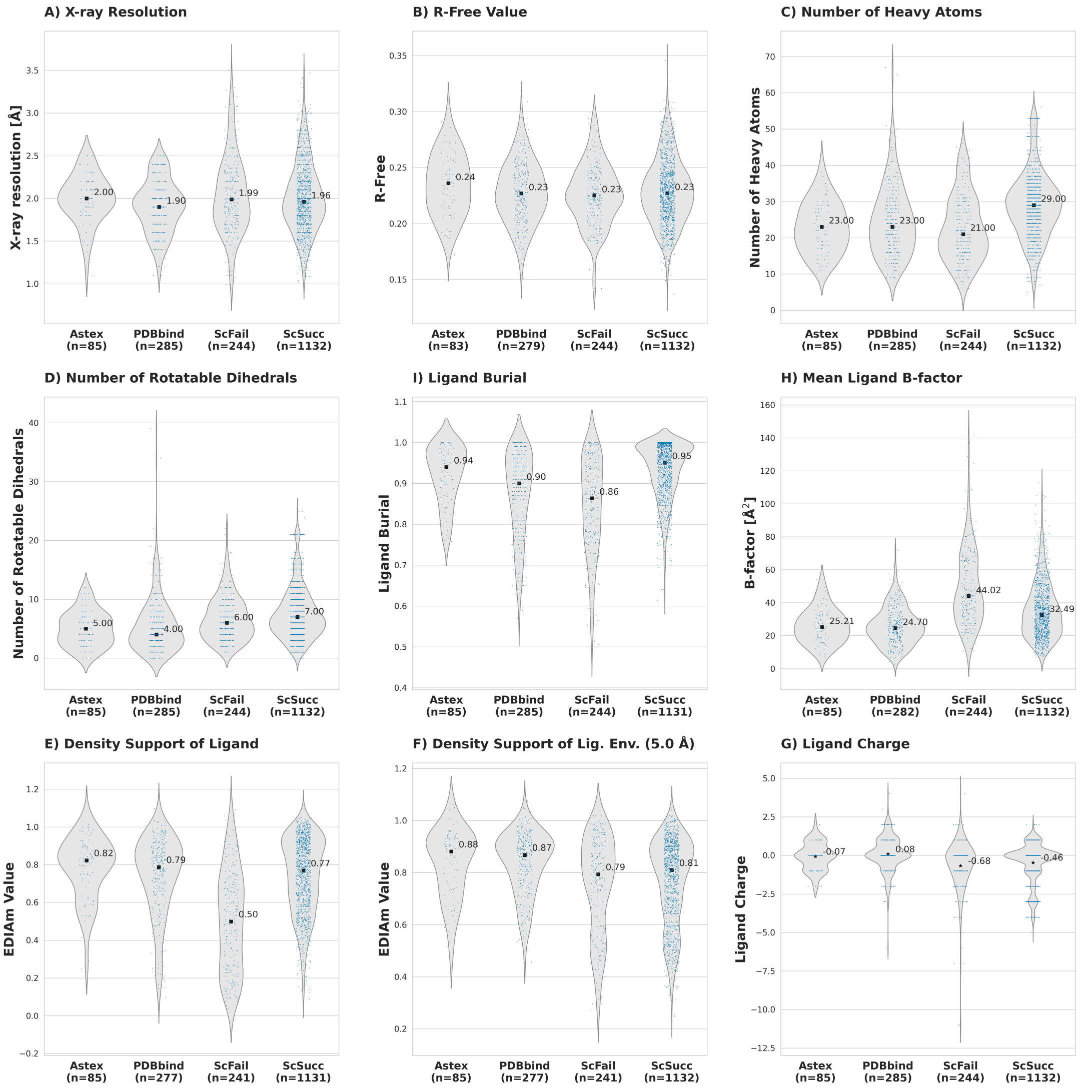

**Figure S5.** Characteristics of Astex and PDBbind benchmark sets compared to RNP cases that are always scoring failures (ScFail) or always scoring successes (ScSucc). Median/mean values are indicated by black squares/stars, respectively, and annotated. On average, scoring failures have smaller and less flexible ligands, which are more solvent exposed, have a higher temperature factor, are more charged, and more ill-defined by the electron density than scoring successes.

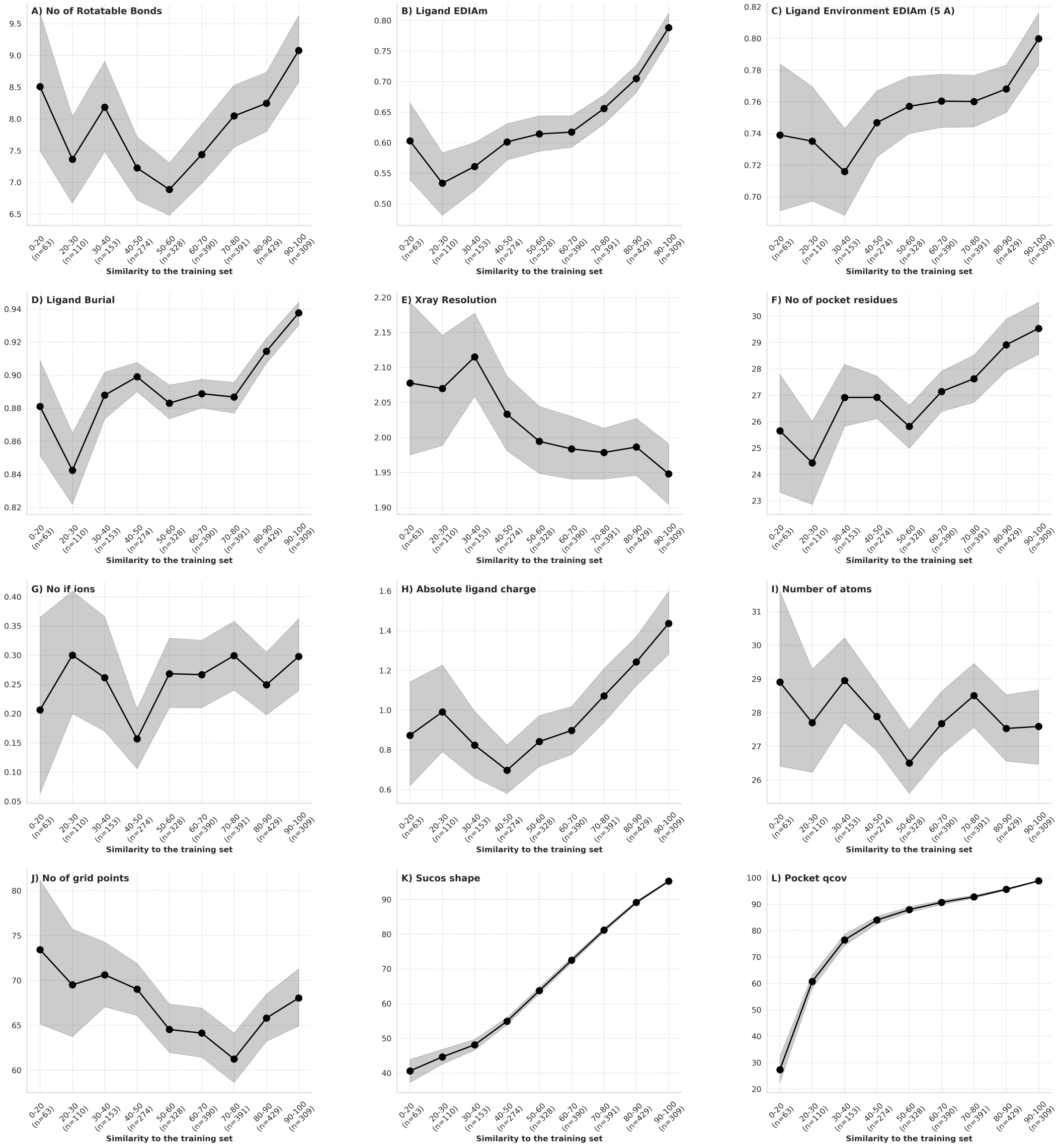

**Figure S6.** Properties of RNP complexes as a function of similarity. Shaded regions correspond to the 95% confidence interval, calculated from 1,000 bootstrap samples for each bin as done in Ref.[1].

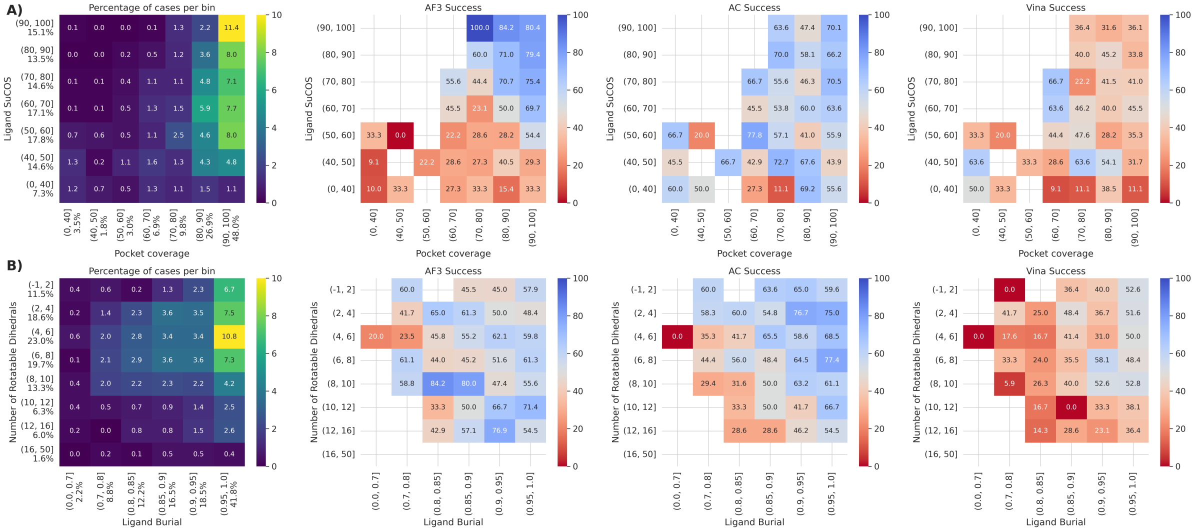

**Figure S7.** Success rates of AF3, AC, and Vina on RNP-F common subset (854 cases) as a function of two properties. The first plot shows the percentage of cases in each bin for the plots on the respective line. A) Success rates as a function of pocket and ligand similarity. B) Success rates as a function of ligand burial and number of rotatable dihedrals.

### References

1. Škrinjar, P., Eberhardt, J., Tauriello, G., Schwede, T. & Durairaj, J. Have protein-ligand cofolding methods moved beyond memorisation? *bioRxiv* 2025.02.03.636309, DOI: https://doi.org/10.1101/2025.02.03.636309 (2025).

2. Daina, A. & Zoete, V. Rethinking molecular beauty in the deep learning era. *bioRxiv* DOI: https://doi.org/10.64898/2025.12.03.692079 (2025).
